# Improved Validation of Protein Interactions using Bicistronic BiFC (Bi2FC)

**DOI:** 10.1101/2024.06.28.601222

**Authors:** Prakash Sivakumar, Vijayaraj Vaishnavi, Kothuri Gayatri, Gayathri R Satheesh, Imran Siddiqi

**Affiliations:** - CSIR–Centre for Cellular and Molecular Biology; Uppal Road, Hyderabad 500007, India; - Academy of Scientific and Innovative Research (AcSIR), Ghaziabad - 201002, India

**Keywords:** BiFC, transient expression, fluorescent protein, P2A, translational reporters

## Abstract

Refolding based Bimolecular Fluorescence Complementation (BiFC) has emerged as an important in vivo technique to identify protein interactions. Significant improvements have been made to enhance the detection capacities of BiFC, however less attention has been paid to the detection of expression levels of proteins. Here we demonstrate development and validation of an improved method to identify protein interactions that incorporates an expression control based on bicistronic expression of the protein of interest and a fluorescent protein separated by a self-cleaving peptide. This method gives robust identification of positive interactions and more reliably identifies absence of interactions. We also show an earlier identified non-interacting pair in yeast two-hybrid (Y2H) to be interacting in vivo.

**Background:** Bimolecular Fluorescence Complementation (BiFC) is a well-utilized method to detect protein interactions in vivo (Kodama and Hu 2012). It is dependent on the refolding of a fluorescent protein split into two parts. The proximity between the two parts leads to functional refolding of the protein and a corresponding increase in the fluorescence intensity. Proteins of interest can be tagged to either part of the split fluorescent protein and be expressed together in a single cell. The physical proximity provided due to interactions between the proteins brings the two parts of the split fluorescent protein closer leading to the formation of a functional fluorescent complex. BiFC has at times been used as the sole technique to identify protein interactions *in vivo* or else as a step following identification of candidate positives by yeast two-hybrid (Y2H). In order to identify the interactions reliably it is also important to validate the gene expression status of BiFC elements (Kudla and Bock 2016). However, systems to visualize the interactions and expression status together are not available for plants.

**Results:** We generated and validated a vector system, called Bi2FC (Bicistronic BiFC) vector system for verifying protein interactions in plants. Bicistronic expression of proteins of interest fused with the two different BiFC components and a fluorescent protein, separated by a self-cleaving peptide identified the expression status of the fusion, while simultaneously identifying positive and negative interactions. As a test, we identified interaction between MPK8 and GRF4 using our system, which were previously identified to not show interaction via Y2H.

**Conclusion:** We described an improved BiFC method to current BiFC to identify the expression of individual proteins tagged with split fluorescent protein as well as to monitor their interaction in a plant cell. This method incorporates additional controls that express only the split fragments together with a vector control, while also providing a set of validated positive and negative controls. Using our method, we also show a previously identified non-interacting pair in yeast two-hybrid (Y2H) to be interacting, emphasizing the need for improved detection and validations.

## Introduction

Bimolecular Fluorescence Complementation (BiFC) is a widely used method to test for protein-protein interactions in vivo (Kodama and Hu 2012). In the BiFC system, a fluorescent protein (FP) is split into two parts as N-terminus and C-terminus (Hu et al. 2002). The N-terminus part of FP is fused with one of the interacting partners, while the other interacting partner is fused to the C-terminus part of FP. When the fusion proteins are made to express in the same cell, in the event of interaction between putative interacting partners, the proximity of both FP fragments promotes association and complete folding of the FP leading to a concomitant increase in the fluorescence signal. In 2016 Kudla and Bock presented community guidelines for BiFC experiments, particularly in plants (Kudla and Bock 2016). This includes the usage of proper controls, the establishment of expression of each split FP fusion in a given cell, quantification, and the usage of alternate in vivo techniques to further validate the interactions. In order to assess the extent to which best practices for BiFC in plants as suggested by (Kudla and Bock 2016) are being followed, we did a systematic sampling of freely accessible PubMed articles that used BiFC in plants from 2021 to 2023. The result of our survey report (Supplementary Table 1) showed that 109 out of 151 articles used BiFC as the only in vivo method to study protein interactions. This suggests the wide acceptability of BiFC in the plant community, as a sufficient standard for in vivo protein identification. Here we demonstrate the generation of a vector system that satisfies one of the guidelines provided by Kudla and Bock which emphasizes the need to establish expression status of BiFC components. The approach uses a **Bi**cistronic **Bi**FC (Bi2FC) system of vectors which provides a readout for the translation of individual split FP fusions in the same cell, and also provides N/C terminus split FP expressible vector controls, in addition to a set of validated positive and negative controls. The system provides a robust method enabling faithful identification of positive as well as negative interactions in plants.

## Results and Discussion

### Bicistronic detection of BiFC components

Fluorescent proteins such as Yellow Fluorescent Protein (YFP) and its variant Venus are extensively used in plants as the split fluorescent proteins for BiFC. Around 155th aa position of YFP protein acts as a split site. (1 - 155 aa) and (156 - 238 aa) polypeptides of YFP are called nYFP and cYFP respectively (Jia et al. 2023). Coding sequences of proteins of interest (POI) to study interactions can be fused in frame with coding sequences of nYFP or cYFP. The nYFP tagged protein is one component of the BiFC while the cYFP tagged protein is the other component. Commonly, the two components are expressed from independent vectors, but single vectors expressing both components have also been used (Grefen and Blatt 2012). Co-expression and association of the split FP fusions in a single cell leads to the refolding of nYFP and cYFP to form a functional complex if the proteins of interest interact. One of the golden rules for BiFC suggested by Kudla and Bock (2016) is that the protein expression status of all the split FP fusions should be tested, including the negative controls.

In plants, BiFC has been performed in Nicotiana, Arabidopsis, onion, and other species. Based on a survey, 64% of the articles used Nicotiana plants in BiFC, with rice and Arabidopsis following closely behind (Supplementary Table 1). The prominence of Nicotiana in BiFC is due to the ease of transient transformation assays of split FP fusions into Nicotiana via Agrobacterium. In a typical BiFC experiment, vectors harboring nYFP: POI1 and cYFP: POI2 are independently transformed into Agrobacterium. Both the Agrobacterium strains are mixed in equimolar concentration and co-infiltrated into the Nicotiana leaves. Agrobacterium-mediated transformation of transfer DNA (T-DNA) containing the split FP fusions is the first step in BiFC. However, T-DNA transformation and corresponding expression of the construct can also show bias in efficiency (Kim et al. 2007). Also, the co-transformation frequency of different T-DNAs was experimentally identified to be less than 30 % (De Buck et al. 2009). To rectify this problem some BiFC vectors are available to report the transformation status in a given cell (Grefen and Blatt 2012). However, those vectors do not indicate the expression status of split FP fusions under study.

A useful system would be one which reports both the transcription and translational status of split FP fusions in a given cell. This can be achieved by creating a bicistronic fusion with spectrally distinct fluorescent proteins other than the split fluorescent protein. Direct visualization of the expression eliminates the need to identify the transformation of T-DNAs. Internal Ribosomal Entry Sites (IRES) (Wong et al. 2008) and self-cleaving peptides such as T2A and P2A (Khosla et al. 2020) from viruses have been proven to be useful bicistronic mediators in various systems. IRES is an alternate entry site for the ribosome other than the canonical 5’-7 methyl guanosine cap (Hinnebusch and Lorsch 2012). However, IRES system acts as a transcriptional reporter rather than a translational reporter. Since a translational reporter can report the transformation of T-DNA as well as transcription and translation of the split FP fusion, we decided to use P2A, as a self-cleaving peptide for the generation of a bicistronic BiFC system.

### Spectrally distinct fluorescent protein reporters for BiFC parts

BiFC works based on the reconstitution of two parts of a fluorescent protein. YFP or Venus fluorescent protein as split fluorescent protein are largely used and other fluorescent proteins like Cyan Fluorescent protein (CFP), mRFP, and mCherry have also been used as BiFC split protein (Jia et al. 2023). Two spectrally distinct fluorescent proteins from the one used as split FP are required for the development of translational reporters in the BiFC system, one to report the expression nYFP:POI1 and another to report the cYFP:POI2. Another issue that has to be taken into consideration while designing reporters for translation in the BiFC system for plants is the strong auto fluorescence of chloroplasts around 650-700 nm (Donaldson 2020). Here, YFP / Venus seems to be an optimal choice for BiFC systems due to the 515 nm excitation maxima (ex) and 528 nm emission maxima (em) falling approximately in the middle of the visible spectrum, providing choices to select FPs in the blue and red range. Hence, it would be useful to have bright fluorescent protein for one compartment around 450 nm and another around 600 nm. Therefore, we chose mTagBFP2 with ex-399 nm and em-452 nm and mCherry with ex −587 nm and em - 610 nm for the generation of bicistronic BiFC.

### Development of Bi2FC vectors

We used a 35S promoter with omega translational enhancer sequences to drive the BiFC components in an Agrobacterium binary vector. Under the regulatory control of pro35S: omega we cloned the N-terminus part of Venus until 172 aa, followed by a gateway cassette and P2A:3xNLS:mCherry fusion. This vector has been named as pBi2FC-N-GW-P2A:mCherry. Similarly, we built pro35S:omega: C-terminus part of Venus (155 - 230 aa) followed by a gateway cassette and further connected by the presence of P2A:3xNLS:mTagBFP2. This vector has been named pBi2FC-C-GW-P2A:mTagBFP2. We also cloned a short linker sequence MGGAGM via gateway recombination into both the Bi2FC vectors and made the controls as pBi2FC-N-MGGAGM-P2A:mCherry and pBi2FC-C-MGGAGM-P2A:mTagBFP2. The putative interactors POI1 and POI2 without stop codons can be cloned into the Bi2FC vectors via Gateway recombination to get pBi2FC-N-POI1-P2A:mCherry and pBi2FC-C-POI2-P2A:mTagBFP2 respectively (Fig. 1).

**Fig. 1:**
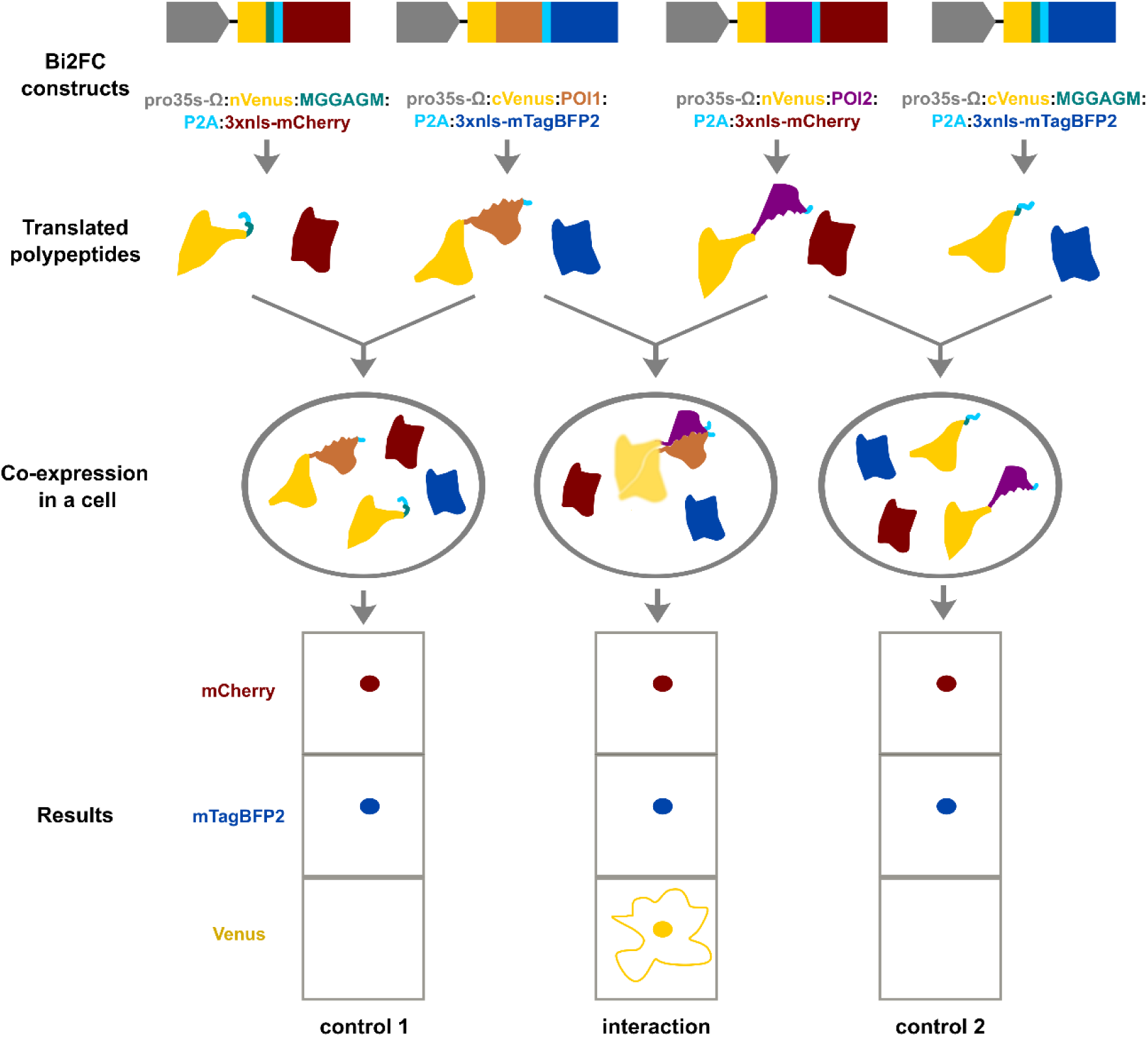
Schematic representation of Bi2FC technique. The Bi2FC constructs are represented as blocks and with corresponding color coded texts. When the Bi2FC constructs are made to express, they produce two polypeptides due to P2A sequence, and corresponding products are colored the same as that of constructs. Co-expression of two constructs in a cell as indicated by connecting arrows leads to the production of 4 polypeptides. The expected result of controls and positive interactions is indicated.

We provide constructs which express only the N or C terminus of split FP separated by a nuclear localized mCherry and mTagBFP2 respectively. It is important to assess the expression status of vector controls and their potential to non-specifically interact with proteins of interest under study. In order to identify protein interactions using the Bi2FC method one should test the following combinations. The combinations (pBi2FC-N-POI1-P2A:mCherry + pBi2FC-C-MGGAGM-P2A:mTagBFP2) and (pBi2FC-C-POI2-P2A:mTagBFP2 + pBi2FC-N-MGGAGM-P2A:mCherry) test for non-specific fluorescence arising due to an interaction between split FP sub fragment and proteins of interest. Combination of pBi2FC-N-POI1-P2A:mCherry + pBi2FC-C-POI2-P2A:mTagBFP2 tests for the interaction between POI1 and POI2. If two Bi2FC constructs are co-infiltrated and the nucleus fluoresces with both mCherry and mTagBFP2, it is indicative of transformation of both the vectors into the same cell and transcription and translation of split FP fusions in that cell (Fig. 1). An additional signal in the Venus channel indicates an interaction between POI1 and POI2 (Fig. 1). The above features of the BiFC vector system which we name Bi2FC, report the transformation of T - DNA, transcription of split FP fusions, and their translation.

### Survey of current BiFC practices and comparison of Bi2FC with existing methods

In order to understand the current BiFC practices and the improvements Bi2FC offers, we used Pubmed’s advanced searching tool to search for BiFC keywords and restricted the search to free articles published from 2021 to 2023. From those articles we categorized the article content pertaining only to BiFC experiments in the context of plants. Nicotiana leaves based transient assay for BiFC in plants is most widely used with 64 % of the articles using it, followed by the use of Arabidopsis protoplasts, rice protoplasts, onion peels and other plants (Supplementary Table 1). Kudla and Bock (2016) suggest the usage of a truly independent in vivo method to reconfirm the protein-protein interactions along with BiFC. However, 109 articles out of 151 used BiFC as the only in vivo technique to verify protein interactions (Fig. 2A). This suggests the predominant reliance on BiFC as the standard for identifying protein interaction identifications. The remaining 42 articles validated interactions using BiFC along with other techniques like co-immunoprecipitation and split luciferase (Fig. 2A).

**Fig. 2:**
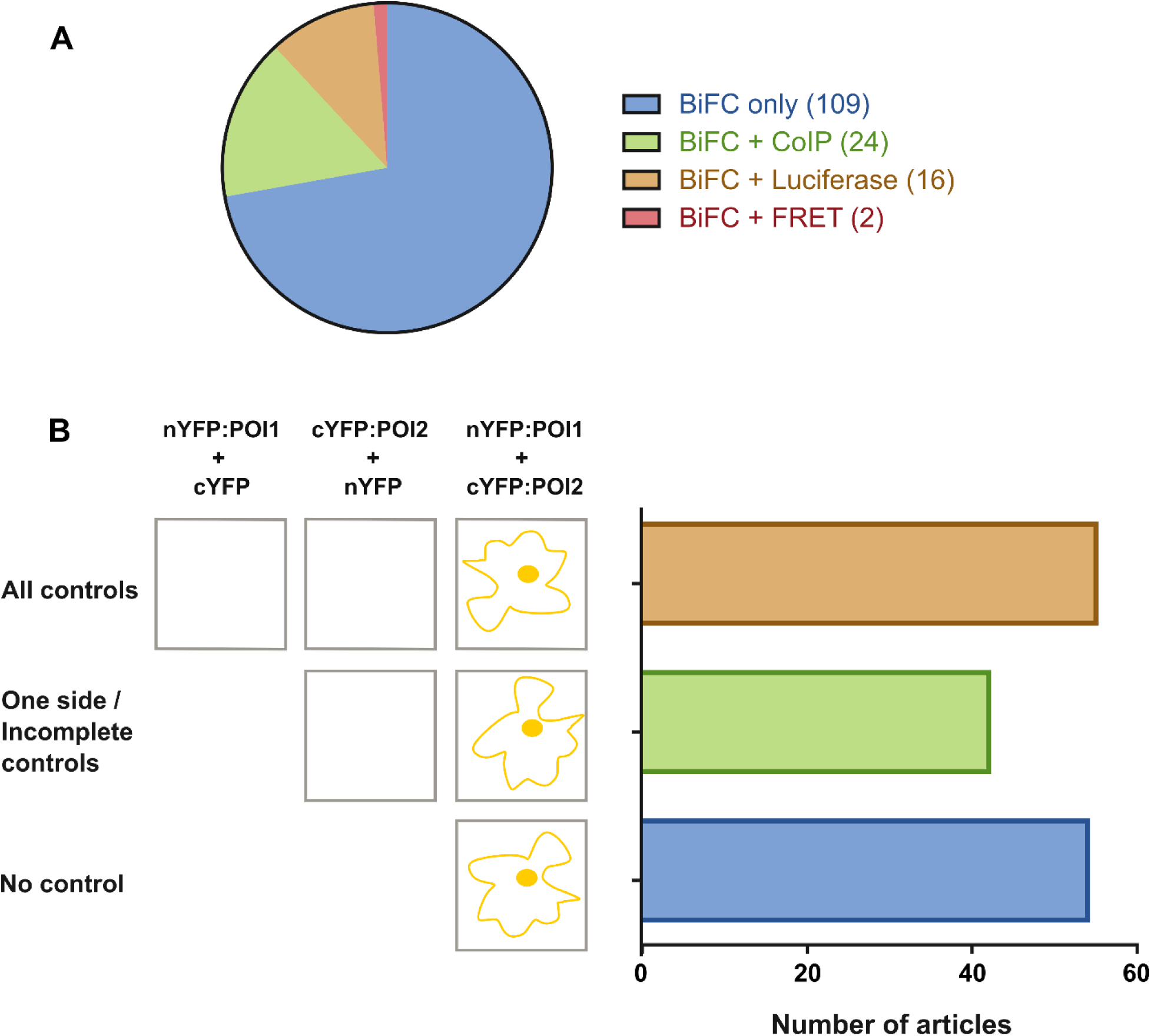
Survey results on BiFC experiments in plants. A) Pie chart of articles which used other in vivo techniques along with BiFC. B) Schematic diagram explaining the usage of controls in BiFC from surveyed publications and their proportion as bar chart.

The widespread reliance on BiFC prompted us to examine the usage of controls in BiFC. A negative control in BiFC tests for non-specific fluorescence between nYFP:POI1 and cYFP. Likewise, nYFP and cYFP:POI2 should also be tested for negative interaction. This ensures the detection of non-specific interactions between the POIs being tested and the components of BiFC. We noted that often the controls which express only nYFP or cYFP were not validated to be expressed. Testing the expression status of the non-fused fragments of YFP in a control is also equally important. Moreover, the usage of starting vectors as either nYFP or cYFP expressing controls is being practiced. The faithful expression of nYFP or cYFP can be affected by the presence of multicloning sites derived cryptic peptides. The issue becomes potentially significant when gateway vectors are used. Gateway components like ccdB, recombination sites and other elements can create cryptic polypeptides which can affect the formation of non-specific interactions.

Of the surveyed articles, 36% of them tested both the n/cYFP:POIs with complement fragment of YFP and 28% of articles either used unclear and ambiguous controls or tested only one of the POIs for negative interaction. Surprisingly, the remaining 36% of articles did not provide any control (Fig. 2B). We reiterate the need to include controls that can express only the YFP fragments as vector control in the BiFC experiments. The Bi2FC system would offer a straightforward solution to the problem of decreased use of negative controls, by having streamlined n/cVenus expressible controls and translational reporter to assess the expression status of the BiFC components (Table 1).

**Table 1:** Comparison of Bi2FC with other methods.

| Property | BiFC | 2in1 BiFC | Bi2FC |
| --- | --- | --- | --- |
| Detection of T-DNA transformation status | No | Yes | Yes |
| Detection of expression of BiFC elements | No | No | Yes |
| Two vector system | Yes | No | Yes |
| Validated split fluorescent protein expressible vector controls | No | No | Yes |

### Validation of Bi2FC system

To test our system, we searched for a protein that has a set of studied positive and negative interactions. We found GRF4, one of the 14-3-3 proteins from Arabidopsis to meet these requirements along with its MAP kinase interactors (Yu et al. 2023). We cloned positive interactors of GRF4, like MPK5, and MPK6 as well as the negative interactors MPK8 and MPK12 into Bi2FC vectors. All the four mentioned MPKs were cloned into Bi2FC vector as pBi2FC-N-MPKx-P2A:mCherry, while GRF4 was cloned as pBi2FC-C-GRF4-P2A:mTagBFP2 vector. Agrobacterium carrying split FP expressible vectors, pBi2FC-N-MPK6-P2A:mCherry and pBi2FC-C-GRF4-P2A:mTagBFP2 were independently infiltrated into Nicotiana leaves to examine the nature of reporter fluorescence with and without fusing with protein of interest in mTagBFP2, Venus and mCherry channels. Irrespective of fusion, constructs with either mTagBFP2 or mCherry as reporter fluoresced in the nucleus of corresponding channels (Supplementary Fig. 1). To test for non-specific interactions, we co-infiltrated Nicotiana with pBi2FC-C-MGGAGM-P2A:mTagBFP2 and pBi2FC-N-MPKx-mCherry and independently co-infiltrated Nicotiana with pBi2FC-C-GRF4-P2A:mTagBFP2 and pBi2FC-N-MGGAGM-P2A:mCherry (Figs. 3,4; Supplementary Fig. 1). As anticipated, we found that mTagBFP2 and mCherry expressed in the nucleus. However, we also observed a non-specific signal in the Venus channel, confined to the nucleus only in coinfiltrated cells. A previous study (Gookin and Assmann 2014) revealed non-specific assembly of split FPs derived from almost all the split sites tested with varying levels of background. We detected non-specific signal only in the nucleus. This could be due to uncleaved polypeptide formed by ribosomal read through of P2A sequence (Liu et al. 2017; Luke and Ryan 2018) being localized to nucleus and nuclear microenvironment enhancing non-specific association of split FPs. Since, MPKs and GRF4 have published interactions in the cytoplasm, we restricted ourselves to the identification of interactions only in the cytoplasm.

**Fig. 3:**
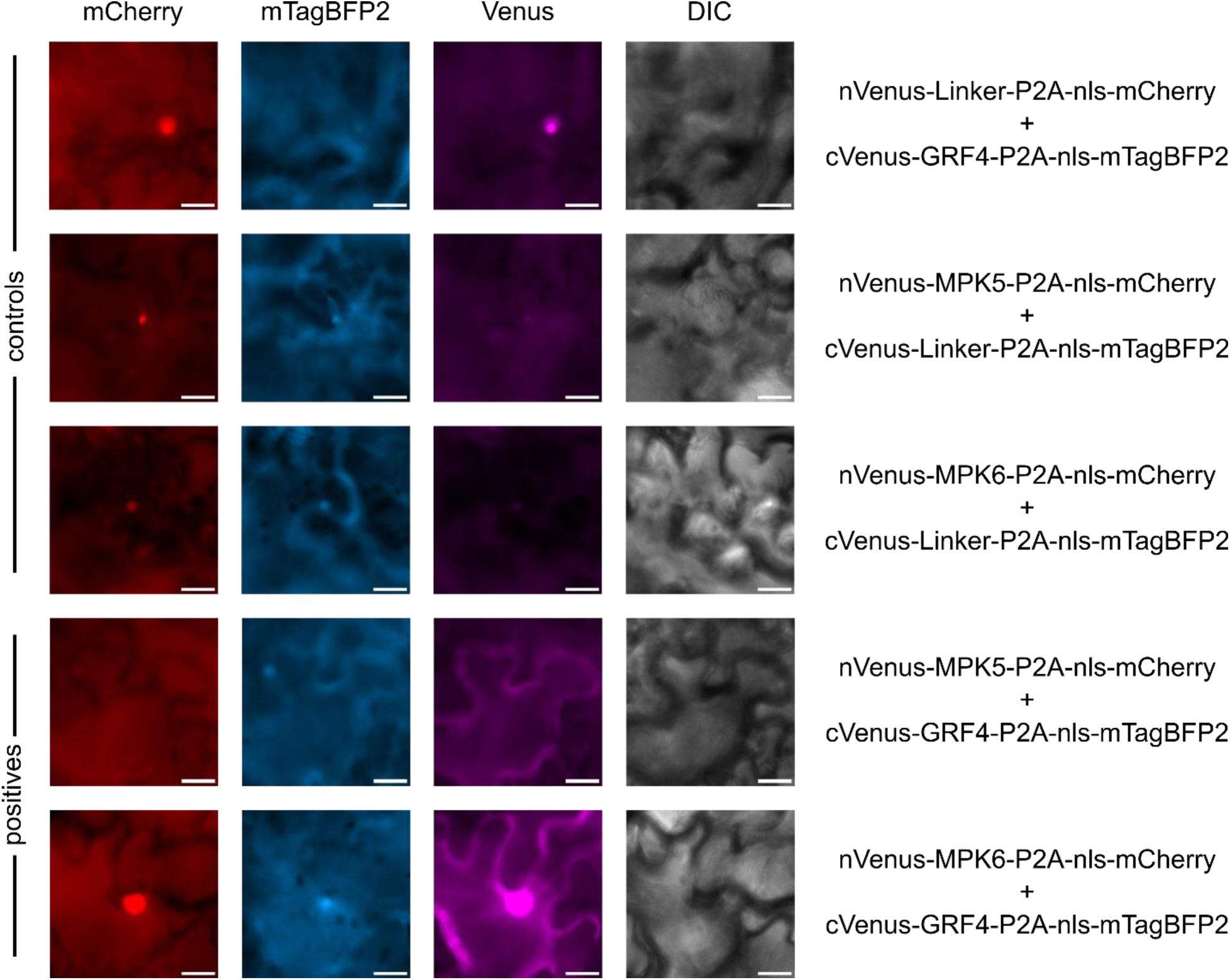
Testing positive interactions using Bi2FC. The controls showed nuclear signal in all three channels as mentioned in text. MPK5 and MPK6 with GRF4 mentioned as positives provided nuclear signal in mCherry and mTagBFP2 channels suggesting the expression of both the split FP fusions, while the irregular shaped cell fluorescing in the Venus channel suggest interactions. Scale - 20 µm.

**Fig. 4:**
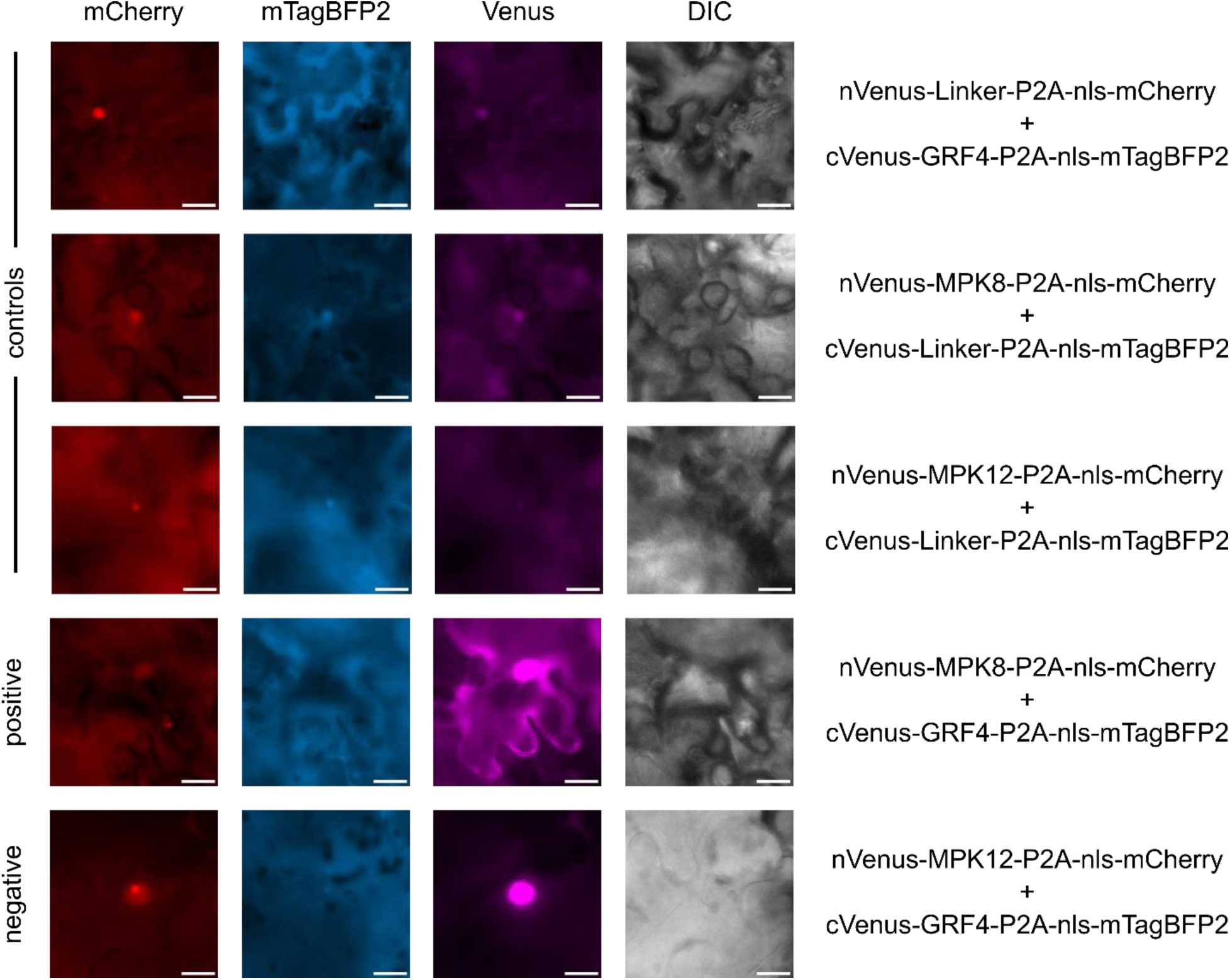
Testing negative interactions using Bi2FC. The controls showed nuclear signal in all three channels as mentioned in the text. Both MPK8 and MPK12 with GRF4 expressed in the shown cell as evidenced by the nuclear fluorescence of mCherry and mTagBFP2. MPK8 with GRF4 is mentioned as positive shown interaction in the irregular cell shape evidenced via Venus fluorescence, while MPK12 with GRF4 mentioned as negative shown no interaction in the cytosol of Nicotiana. Scale - 20 µm.

We reconfirmed the interactions of MPK5 and MPK6 with GRF4 using the Bi2FC system (Fig. 3). We note the varying expression status of different fusion proteins. For example, MPK6 had better expression than MPK5 (Fig. 3). The expression bias could pose challenges in delivering stoichiometrically comparable amounts of proteins to a given cell. MPK8 and MPK12 were negative interactors with GRF4 as per Y2H experiments (Yu et al. 2023). Hence the authors did not test for their interaction status in vivo. We observed no interaction between MPK12 and GRF4 and identified an interaction between MPK8 and GRF4 using the Bi2FC system (Fig. 4). Identification of a MPK8-GRF4 interaction points to the usefulness of employing independent methods for testing and validating interactions in planta.

### Limitations and potential improvements of Bi2FC

One limitation of the Bi2FC system is its inability to identify nuclear interactors, due to non-specific fluorescence in the nucleus. Arabidopsis has a predicted nuclear proteome of 10 – 20 % of proteins (Narula et al. 2013), and there are known nuclear proteins showing interactions in the cytosol (Böttner et al. 2009; Yang et al. 2019; Maika et al. 2023). Hence, the limitation does not preclude the usability of the system in such scenarios and users can also target proteins of interest to the cytosol by fusing with a nuclear exit signal (NES) to test for interactions in the case of nuclear proteins.

## Materials and Methods

### Generation of vectors and constructs

The mTagBFP2 (gift from Fabio Pasin) and mCherry were amplified with primers 1202 and 1203, and 1202 and 172 to produce P2A:3xNLS:mTagBFP2 and P2A:3xNLS:mCherry with XhoI and SacI sites are cloned into pDEST-VYCE(R)GW and pDEST-VYNE(R)GW respectively to make the intermediary vector, and were digested with SacI and XbaI and cloned into pGWB502-omega to make the pBi2FC-C-GW-P2A:mTagBFP2 and pBi2FC-N-GW-P2A:mCherry respectively.

The clones of MPK5, MPK6, MPK8, MPK12 and GRF4 without stop codon were amplified from Arabidopsis cDNA made from the unopened flowers using primers mentioned in Supplementary Table 2 with NotI and AscI sites and cloned into Gateway entry vector pENTR-D-TOPO digested with NotI and AscI. The MGGAGM linker sequence was ordered as oligo (1218 and 1219), annealed and cloned into pCONR221 via BP cloning. The expression constructs were made via Gateway LR recombination into Bi2FC vectors. All the sequence files are submitted as a genbank file (Supplementary Text). The plasmids discussed in the study are made available via Addgene IDs 216136 to 216146 (https://www.addgene.org/browse/article/28244171/).

### Agrobacterium mediated transformation and imaging

All the expression clones were transformed into the GV3101 strain of Agrobacterium. The strains were infiltrated using 0.8 OD600 of each of the desired combinations mixed equally in the MES buffer. The adaxial side of the 1 month old Nicotiana leaves were infiltrated. The samples were examined after 72 - 80 hrs post infiltration by inverting the leaves to show the adaxial side and imaged using AxioImager2 fluorescence microscope fitted with mTagBFP2, Venus, and mCherry filter cubes. The filter cubes have the following specifications of Exciter Band Pass (Ex-BP) wavelength range in nm, Beam Split (BS) wavelength in nm, and Emitter Band Pass (Em-BP) range in nm; mTagBFP2 (Ex-BP 335-383 nm : BS 395 nm : Em-BP 420-470 nm), Venus (Ex-BP 490-510 nm : BS 515 nm : Em-BP 520-550 nm), and mCherry (Ex-BP 533-558 nm : BS 570 nm : Em-BP 570-640 nm). Images were captured under false colour, analyzed using ImageJ, and assembled in Inkscape.

## Author contributions

PS, VV, and IS conceptualized the project. PS, VV, KG, and, GRS collected the data. PS wrote the first draft. PS and VV made the figures and illustrations. PS, VV, KG and, IS edited the manuscript. IS acquired funding and managed the project.

## Supplementary Materials

Supplementary materials for this article are available and provided separately.

Supplementary Table 1 – BiFC survey data

Supplementary Table 2 – Primers list

Supplementary Figure 1 – Checking for cross excitations in various channels

Supplementary Text – Genbank files of Bi2FC constructs used in this study.

## Acknowledgement

Authors thank Fabio Pasin for the gift of mTagBFP2, CCMB microscopy facility, Mekala Saikiran for plant growth room maintenance, and Priyanka Pant for comments on the manuscript. PS is a recipient of CSIR fellowship. IS acknowledges a JC Bose fellowship from the Department of Science and Technology and a Senior Scientist fellowship from the Indian National Science Academy. This work was supported by a CoE grant to IS from the Department of Biotechnology and a grant from CSIR.

## Conflicts of interest

The authors declare no conflicts of interest.

---

**Supplementary Figure 1:**
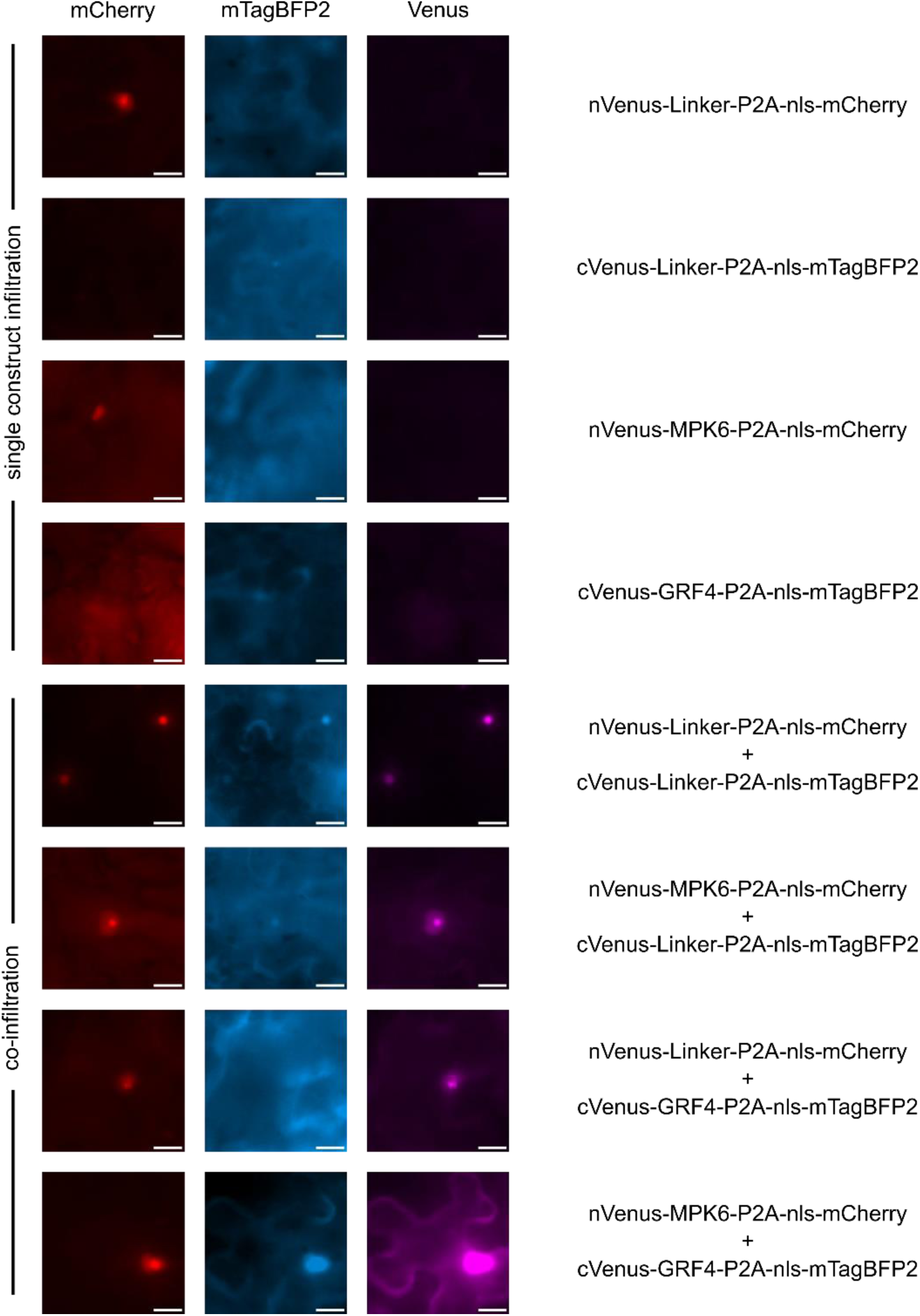
Validation of Bi2FC and test of signal specificity. Agrobacterium mediated infiltration of single vector constructs as mentioned gave nuclear signal only in the corresponding channel. When nVenus and cVenus vector constructs were co-infiltrated, apart from the corresponding reporter specific fluorescence, a non-specific Venus signal was observed in the nucleus. nVenus-MPK6-P2A-nls-mCherry plus cVenus-GRF4-P2A-nls-mTagBFP2 combination showed Venus fluorescence in the cytoplasm along the cell outline. Scale – 20 µm.

## Notes

https://www.addgene.org/browse/article/28244171/

### Competing Interest Statement

The authors have declared no competing interest.

