## Supplementary Table 2 for "Improved Validation of Protein Interactions using Bicistronic BiFC (Bi2FC)"

| **Primer Name** | **Primer Full Name** | **5'-3' Single stranded sequence** |
| --- | --- | --- |
| 1218 | 1218_attB1_OL_NO_STOP_F | GGGGACAAGTTTGTACAAAAAAGCAGGCTTAATGGGTGGTGCAGGAATGAACCCAGCTTTCTTGTACAAAGTGGTCCCC |
| 1219 | 1219_attB2_OL_NO_STOP_R | GGGGACCACTTTGTACAAGAAAGCTGGGTTCATTCCTGCACCACCCATTAAGCCTGCTTTTTTGTACAAACTTGTCCCC |
| 1202 | 1202_Glowing_Protein_P2A_3xNLS_XhoI_F | TACAAAGTGGTTCTCGAGgcaaccaactttagcctgctcaagcaagcaggagatgttgaggaaaatcctggcccccctAAGAAAAAGAGAAAGGTTccGAAAAAGAAAAGAAAGGTGccaAAAAAAAAGAGAAAGGTAATGGTGAGCAAGGGCGAGGA |
| 1203 | 1203_SacI_mTagBFP2_R | GTTTGAACGATCGGGGAAATTCGAGCTCTTAATTAAGTTTGTGCCCCAGTTT |
| 172 | 172_mCherry_SacI_R | GTTTGAACGATCGGGGAAATTCGAGCTCTTACTTGTACAGCTCGTCCATGC |
| 1463 | 1463_GRF4_NotI_F | AGCAGGCTCCGCGGCCGCaATGGCGGCACCACCA |
| 1464 | 1464_GRF4_AscI_R | AAAGCTGGGTCGGCGCGCCtGATCTCCTTCTGTTCTTCAGCAGG |
| 1473 | 1473_MPK5_NotI_F | AGCAGGCTCCGCGGCCGCaATGGCGAAGGAAATTGAATCAGC |
| 1474 | 1474_MPK5_AscI_R | AAAGCTGGGTCGGCGCGCCtAATGCTCGGCAGAGGATTG |
| 1475 | 1475_MPK6_NotI_F | AGCAGGCTCCGCGGCCGCaATGGACGGTGGTTCAGGT |
| 1476 | 1476_MPK6_AscI_R | AAAGCTGGGTCGGCGCGCCtTTGCTGATATTCTGGATTGAAAGCAAG |
| 1479 | 1479_MPK8_NotI_F | AGCAGGCTCCGCGGCCGCaATGGGTGGTGGTGGGAAT |
| 1480 | 1480_MPK8_AscI_R | AAAGCTGGGTCGGCGCGCCtAGAATTGTGAAGAGAAGCAACTTTATCAG |
| 1487 | 1487_MPK12_NotI_F | AGCAGGCTCCGCGGCCGCaATGGATTTAGTGTCTTCAAGAGATACTTTAGG |
| 1488 | 1488_MPK12_AscI_R | AAAGCTGGGTCGGCGCGCCtGTGGTCAGGATTGAATTTGACAGAC |
